# Unravelling the Proteomic Landscape of Imipenem Resistance in *Pseudomonas aeruginosa*: A Comparative Investigation between Clinical and Control Strains

**DOI:** 10.1101/2024.09.20.614093

**Authors:** Marius Eidsaa, Animesh Sharma, Markus Janasch, Alicja Kuch, Anna Skoczyńska, Francesca Di Bartolomeo

**Affiliations:** Department of Biotechnology and Nanomedicine, SINTEF Industry, Trondheim 7034, Norway; Department of Epidemiology and Clinical Microbiology, National Medicines Institute (NMI), Warsaw, Poland; Department of Clinical and Molecular Medicine, Norwegian University of Science and Technology (NTNU), N-7489 Trondheim, Norway; Proteomics and Modomics Experimental Core (PROMEC) at NTNU and the Central Norway Regional Health Authority, Trondheim, Norway

**Keywords:** Antimicrobial Resistance _1_, *Pseudomonas aeruginosa* _2_, Label-free Quantitative Proteomics _3_, antibiotic challenge _4_, imipenem _5_

## Abstract

The increasing prevalence of antimicrobial resistance (AMR) poses a significant challenge to global health, particularly with bacterial pathogens such as *Pseudomonas aeruginosa*, a notorious cause of nosocomial infections. This study focuses on the comparative proteomic analysis of an imipenem-resistant strain of *P. aeruginosa*, a representative of world epidemic clone ST235, and a wildtype control strain, *P. aeruginosa* ATCC 27853, in response to varying concentrations of imipenem. Using label-free quantification (LFQ) and gene ontology (GO) enrichment analyses, we identified significant differences in the proteomic responses between the two strains (data available via ProteomeXchange with identifier PXD055744.). The clinical strain exhibited a stable proteomic profile across the imipenem gradient, suggesting pre-established and efficient resistance mechanisms that do not require extensive reconfiguration under antibiotic pressure. In contrast, the control strain showed a broader, more reactive proteomic response, particularly in proteins associated with membrane transport, stress response, and biofilm formation. Notably, uncharacterized proteins were significantly upregulated in the clinical strain, indicating potential novel resistance mechanisms. These findings highlight the distinct strategies employed by the two strains, with the clinical strain’s stable resistance mechanisms contrasting sharply with the control strain’s reactive approach. The study underscores the importance of further research into the uncharacterized proteins that may play crucial roles in antibiotic resistance, potentially leading to new therapeutic targets in the fight against AMR.

**Highlights:**

1. **Stable resistance mechanisms**: The clinical strain of *Pseudomonas aeruginosa* (ST235) exhibited a stable proteomic profile across varying concentrations of imipenem, indicating pre-established and efficient resistance mechanisms.
2. **Reactive proteomic response**: The control strain (*P. aeruginosa* ATCC 27853) showed a broader and more reactive proteomic response, particularly in proteins related to membrane transport, stress response, and biofilm formation.
3. **Uncharacterized proteins**: Significant upregulation of uncharacterized proteins in the clinical strain suggests potential novel resistance mechanisms that warrant further investigation.

## 1 Introduction

Antimicrobial resistance (AMR) has emerged as a global health crisis, compromising the efficacy of traditional therapies against a diverse array of infections caused by bacteria, parasites, and fungi (Darby *et al*., 2023). This intricate biological phenomenon is primarily fueled by the selective pressure imposed by the widespread and often indiscriminate utilization of antimicrobials, which fosters an environment conducive to the survival and proliferation of resistant strains. The consequent outcome is an escalating prevalence of hard-to-treat infections (Fair and Tor, 2014).

Particularly alarming is the resistance demonstrated by bacterial pathogens. These organisms harness mechanisms such as horizontal gene transfer, mutation and selection, and adaptive resistance, to rapidly evolve and withstand antimicrobial agents (Davies and Davies, 2010). The rise of multi-drug resistant (MDR) and extensively drug-resistant (XDR) bacterial strains adds another layer of complexity to this scenario. These “superbugs” demonstrate resistance against multiple classes of antibiotics, posing serious therapeutic challenges and undermining patient outcomes (Magiorakos *et al*., 2012; Aslam *et al*., 2018).

*Pseudomonas aeruginosa* is an exemplar of such a resilient bacterial species. A notorious instigator of nosocomial infections worldwide, it demonstrates an exceptional ability to resist antibiotic therapies (Breidenstein, de la Fuente-Núñez and Hancock, 2011; Langendonk, Neill and Fothergill, 2021). Given the robust resistance of such strains, treatment often resorts to last-line antibiotics, such as those in the carbapenem class. Imipenem, a member of this class, has a broader antimicrobial spectrum and higher potency than other beta-lactam antibiotics. It operates by interfering with bacterial cell wall synthesis, thus exerting a bactericidal effect. This antibiotic is often seen as a therapeutic beacon against a variety of MDR bacterial infections (Lautenbach *et al*., 2006; Xu *et al*., 2020). However, the growing overreliance on imipenem has inadvertently catalyzed the emergence of imipenem-resistant *P. aeruginosa* strains, escalating the challenges associated with this formidable pathogen. A prime example of its adaptability is the clinical strain P. aeruginosa, recognized as a representative of world epidemic clone of sequence type ST235, owing to its extensive prevalence and heightened resistance to multiple antibiotics.

To effectively counter this rising tide of resistance, we must delve deeper into the molecular mechanisms underpinning the resistance of bacteria to potent antibiotics such as imipenem.

Proteomic analysis, in this regard, can prove instrumental by elucidating antibiotic resistance mechanisms at the protein level, as proteins are the ultimate executors of most cellular functions and interactions (Pérez-Llarena and Bou, 2016). Proteomics can offer invaluable insights into the adaptive responses of bacteria to antibiotic challenges. It achieves this by quantifying changes in protein expression, modifications, and interactions under different conditions. Proteome-wide label-free quantification, a technique frequently employed in proteomic analysis, allows for the comparison of protein abundance between resistant and susceptible strains, or between bacteria subjected to different antibiotic concentrations (Goodyear *et al*., 2023). As for this study, comparative proteomic studies serve as a potent investigative tool for understanding the profound and complex responses of bacteria to antibiotic exposure. They have an unprecedented ability to illuminate the molecular landscape where the dynamic interplay between antibiotics and bacteria unfolds. In the face of antibiotic stress, bacteria respond by expressing an array of proteins that enable them to neutralize the antibiotic’s effect, remove the antibiotic from the bacterial cell, or modify the bacterial cell targets to evade the antibiotic’s impact (Khodadadi *et al*., 2020). Proteomic studies, through their ability to quantitatively analyze the entire complement of proteins expressed under different conditions, enable us to delineate this intricate molecular proteomics pool in detail. The proteomic response of bacteria to antibiotics, which includes the alterations in protein expression, interactions, and modifications, is part of a larger adaptive response of bacteria to survive under antibiotic stress. This response forms the “resistome” of bacteria, a term coined to describe the collection of all the antibiotic resistance genes in a bacterial cell or community (Wright, 2007). While the concept of the resistome is predominantly genomic in nature, the functional output of the resistome is embodied in the proteome. It is the proteins, being the functional units of cells, that execute the instructions coded in the resistome, thereby manifesting the resistance phenotype (Sulaiman and Lam, 2022).

In the present study, we utilize the power of proteomics to dissect the resistome in action, as we analyze how two strains of *P. aeruginosa* -the representative of the world epidemic clone ST235 and the control strain ATCC 27853 - respond to the antibiotic imipenem, scrutinizing these profiles across a spectrum of escalating minimum inhibitory concentrations (MICs) of the antibiotic. This nuanced approach will enable us to trace the dynamic shifts in protein expression and interactions as antibiotic pressure intensifies. Additionally, comparing the two selected strains will yield valuable insights into the proteomic signatures tied to the clinical strain’s heightened resistance. Therefore, this research is primed to enrich our understanding of *P. aeruginosa*’s proteomic responses and the resistance mechanisms to imipenem. These insights may lead us towards potential solutions to the urgent and ongoing crisis of antimicrobial resistance by identifying proteins or pathways that could serve as novel targets for drug development or interventions that could disrupt the functionality of the resistome.

## 2 Materials and Methods

### 2.1 Strains

*Pseudomonas aeruginosa* 6378/09 strain (NMI collection number), hereinafter referred to as clinical strain and *Pseudomonas aeruginosa* ATCC 27853 control (reference) strain, obtained from American Type Culture Collection, were used for proteomic analysis. It is wildtype, imipenem-sensitive strain, hereinafter referred to as control strain. The *P. aeruginosa* clinical strain was isolated from an oncology patient from an abdominal swab. It is imipenem-resistant strain, associated with VIM-2 variant of metallo-beta-lactamase (MBL) and belonged to world epidemic clone ST235, as previously described (Urbanowicz *et al*., 2021). ST235 has been the most frequent MDR *P. aeruginosa* clone globally, associated with VIM/IMP-like MBLs (Oliver *et al*., 2015).

### 2.2 Determination of the minimum inhibitory concentration (MIC)

Minimum inhibitory concentration (MIC) of imipenem was determined for both strains by the broth microdilution method according to standard ISO 20776-1 (ISO 20776-1, 2020). The MIC was defined to be the lowest concentration at which no visible growth (complete inhibition of growth) of bacteria could be observed after incubation for 18±2 hours as recommended by the European Committee on Antimicrobial Susceptibility Testing (EUCAST)(European Committee on Antimicrobial Susceptibility Testing, 2024).

### 2.3 Cell culture and antibiotic challenge

The bacteria were grown in cation-adjusted Mueller-Hinton broth (Becton Dickinson, Sparks, MD, USA) at 37 °C with shaking (120 rpm) to optical density OD_600_ = 1.0. For antibiotic challenge, imipenem concentration of 0.1x MIC, 0.25x MIC, 0.5x MIC and 1x MIC of the particular strain was used in a volume of 5 ml. Bacterial cultures without the antibiotic were considered as untreated controls, **see Table 1**. Challenged bacterial cultures were then incubated with shaking (120 rpm) for 5 hours at 35 °C ± 2 °C. Each experiment was performed three times for both clinical and control strains (3 biological replicates) with two technical replicates for each condition (antibiotic concentration).

**Table 1:**
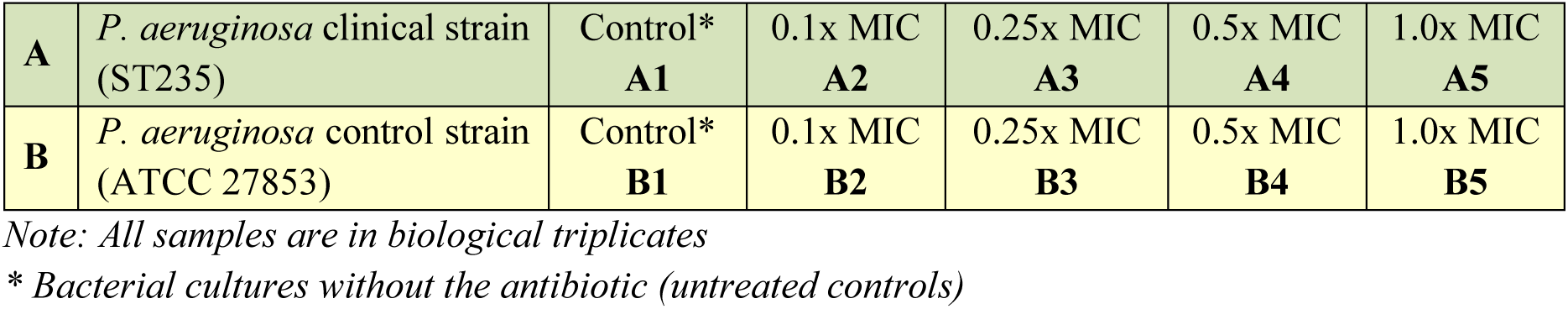
Clinical strain (A) and control strain (B)

After incubation the bacterial cell pellets were harvested in pre-cooled Falcon tubes by centrifugation (5 min at 3,000 × g below 0 °C). Then, the cell pellets were washed twice with ice-cold 1 x PBS (Sigma-Aldrich, Life Science, St. Louis, MO, USA). Finally, cell pellets were collected by centrifugation (5 min at 3,000 × g below 0 °C), snap-frozen, and stored at −80 °C until further testing (Liu *et al*., 2016; Hashemi *et al*., 2019; Jongers *et al*., 2021).

### 2.4 Label-Free Proteomics analyses

Proteins were quantified by processing the timsTOF-pro DDA MS-data using MaxQuant v2.0.3.0 (Tyanova, Temu and Cox, 2016). Namely, the following search parameters were used: enzyme specified as trypsin with a maximum of two missed cleavages allowed; minimum peptide length of 7; acetylation of protein N-terminal, oxidation of methionine, and deamidation of asparagine/glutamine as dynamic post-translational modification. These were imported in MaxQuant which uses m/z and retention time (RT) values to align each run against each other sample with a minute window match-between-run function and 20 mins overall sliding window using a clustering-based technique. These were further queried against the reference-proteome of *Pseudomonas aeruginosa* downloaded from UniProt (UniProt Consortium, 2022) in May 2022 and MaxQuant’s internal contaminants database using Andromeda built into MaxQuant. Both Protein and peptide identifications false discovery rate (FDR) was set to 1%, only unique peptides with high confidence were used for final protein group identification. Peak abundances were extracted by integrating the area under the peak curve. Each protein group abundance was normalized by the total abundance of all identified peptides for each run and protein by calculated median summing all unique peptide-ion abundances for each protein using label-free quantification (LFQ) algorithm (Cox *et al*., 2014) with minimum unique peptide(s) ≥ 1. The mass spectrometry proteomics data have been deposited to the ProteomeXchange Consortium via the PRIDE (Perez-Riverol *et al*., 2022) partner repository with the dataset identifier PXD055744 (Username: Password: XNtCwVbI4AIs).

### 2.5 Data processing and mining

Having obtained the label-free quantification (LFQ) intensity values for both the *Pseudomonas aeruginosa* clinical strain and control strain, across the five different concentrations (as depicted in **Figure 1**), we performed a series of data filtering, processing, and aggregation steps, including removal of reverse hits and potential contaminants to ensure the integrity of our data. In the case of multiple UniProt protein IDs per hits, the one with the highest curation status were used in the downstream analysis when relevant. This led to a total of *N*=3811 proteins in our analysis, accounting for approximately 60% of the chromosomal genes in an average *P. aeruginosa* strain (Mosquera-Rendón *et al*., 2016).

**Figure 1.**
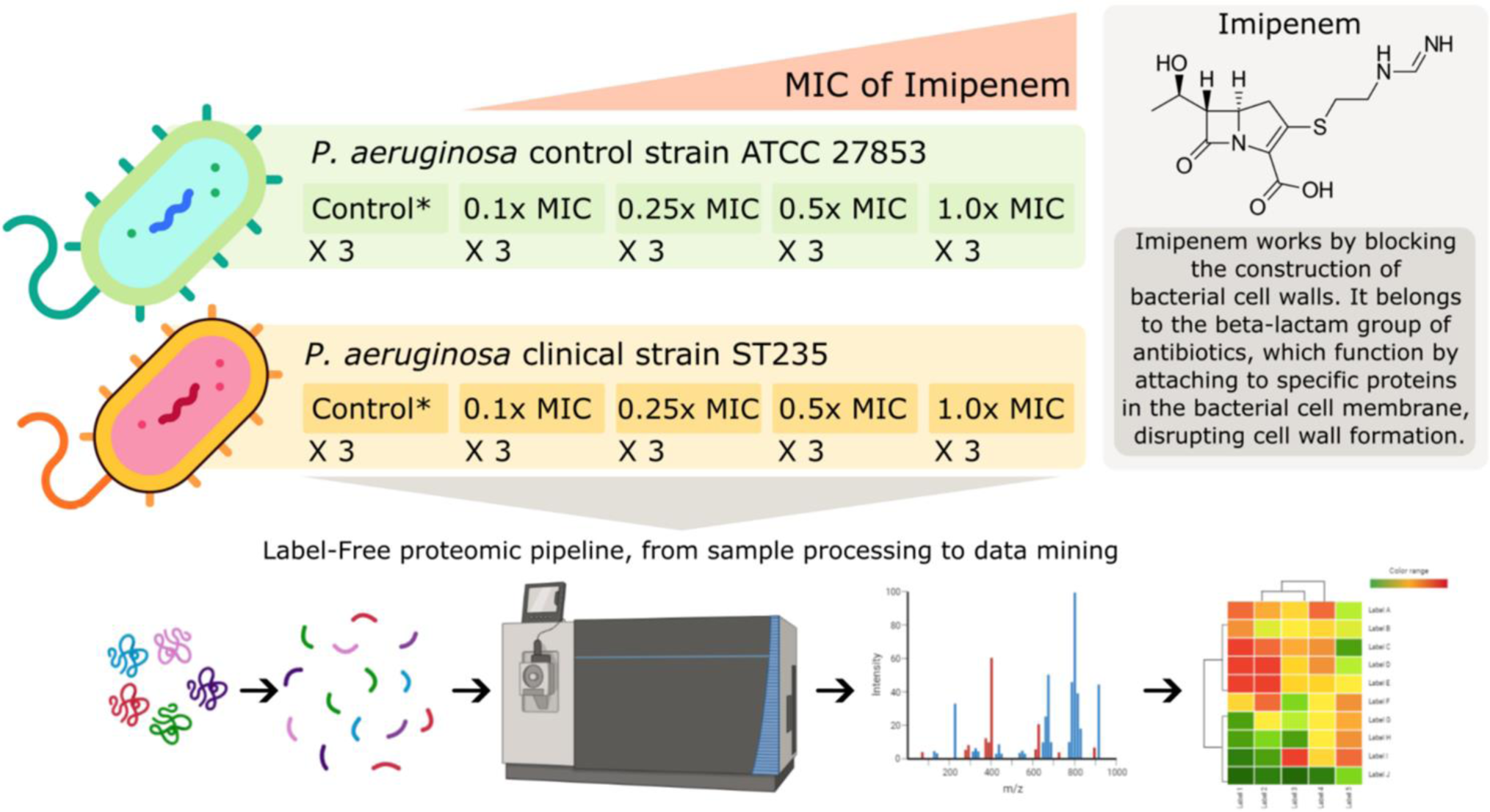
Graphical abstract. Setup and workflow used in the comparative proteomic analysis of Pseudomonas aeruginosa strains under varying concentrations of the antibiotic imipenem. The study involves a P. aeruginosa clinical strain (ST235) and a control strain (P. aeruginosa ATCC 27853), both subjected to increasing concentrations of imipenem based on their respective minimum inhibitory concentrations (MICs) with each condition replicated three times. Imipenem concentrations range from 1.6-16 mg/L and 0.2-2.0 mg/L for the clinical and control strains respectively, corresponding to 0.1x MIC, 0.25x MIC, 0.5x MIC and 1.0x MIC. The untreated control probes (no antibiotic) for each strain are indicated by Control* in the figure. Imipenem, a beta-lactam antibiotic, disrupts bacterial cell wall formation by binding to specific proteins in the bacterial cell membrane. The proteomic pipeline includes sample processing, mass spectrometry for label-free quantification, and data mining, leading to insights into differential protein expression and resistance mechanisms. Part of the figure has been created with BioRender.com (Eidsaa, M. (2024) BioRender.com/w02t614).

For each strain and antibiotic concentration, we had three biological replicates (**Figure 1**). We performed a standard *t*-test using the log2-transformed LFQ (log2LFQ) values for each protein ID to obtain the associated *p*-values. These *p*-values were then corrected using the Benjamini-Hochberg False Discovery Rate (FDR) procedure. This was done for all experiments with non-zero MIC, using the 0.0x MIC as a control for each strain. This means that we compared LFQ values and performed *t*-tests within strains only. Alongside the *t*-test, *p*-value and the FDR values, we used the median value for LFQ intensities where applicable.

When comparing LFQ intensity values between experiments, we primarily used fold change relative to the control probe (no antibiotic) to describe the difference in log2 space, expressed via log2LFQ values. Proteins with LFQ intensity values of zero for both the control probe (no antibiotic) and 1.0x MIC experiments were excluded. Proteins with values of zero in one experiment and non-zero in another were assigned a large, finite fold-change value. When comparing across strains, we used the median log2LFQ fold changes between the strains, as these values were already normalized with respect to the 0.0x MIC control experiment for the respective strains.

Unless otherwise specified, the intra-strain protein LFQ intensity fold-change cutoff was set at 1.5, or approximately +/- 0.5849 after log2 transformation. In the individual protein volcano plots an uncorrected *p*-value cutoff of 0.1 was employed, with values above colored grey and values below colored light red or blue, while FDR values below 0.1 are furthermore indicated by dark red or blue color. These thresholds are not particularly conservative, which is deemed appropriate given the explorative nature of the study. The individual protein volcano plot x-axis and y-axis ranges were fixed to improve visual comparability, and log2 fold changes were visually capped at +/- 5 alongside *p*-values below 1e-5.

To visualize the overall features of the data set, we calculated and used a Principal Component Analysis (PCA), using standard scaling, and a correlation matrix (heatmap) of Spearman’s correlation coefficients between experiments. This provided an overview of the structure and relationships in the data set (**Figure 2**).

**Figure 2:**
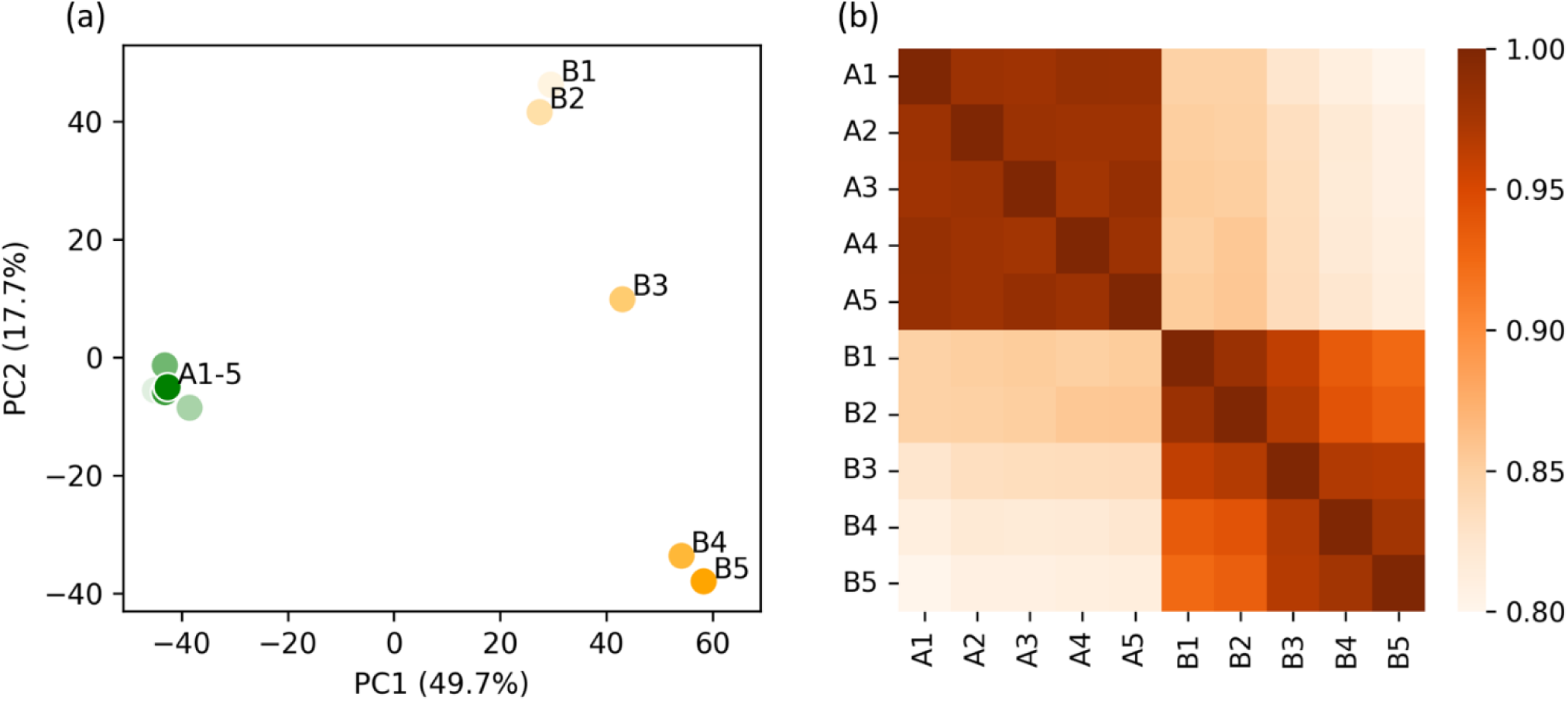
Visualization of the label-free quantification (LFQ) proteomics data sets, illustrating the impact of increasing imipenem concentration on the clinical (A1-5) and control (B1-5) strains. (a) Standard-scaled PCA plot shown for the two most significant principal components, accounting for 49.8% and 17.7% of the total variance. The imipenem-resistant clinical strain forms a compact cluster on the left-hand side, while the control strain exhibits a spread on the right-hand side. This spread shows a clear gradient from low antibiotic concentrations (lighter colors) at the top to high antibiotic concentrations (darker colors) at the bottom. (b) Heatmap of Spearman’s correlation coefficients indicating a similar trend as observed in the PCA plot. It indicates a high overall correlation between the LFQ intensity scores, and particularly high correlations within the clinical strain.

### 2.6 Gene-ontology enrichment analysis

To discern significant systemic alterations in the proteomes within and across strains and antibiotic concentrations, we conducted a Gene Ontology (GO) term enrichment analysis. This method is set-based, meaning that groups of proteins are compared rather than individual proteins. Statistical tests are performed by comparing a subset to the full background set. The GO term enrichment analysis aims at finding enriched functional, process-related, and compartmental annotations associated with each protein in the subset. Collectively, numerous individually insignificant protein-level changes can be linked to statistically significant features on aggregate, cellular level.

The GO term enrichment analysis is agnostic to the subset selection method. Therefore, we employ both FDR-value, uncorrected *p*-values, and no *p*-value thresholds when selecting proteins, in conjunction with fold-change thresholds, at various parts of the analysis. We performed statistical tests based on hypergeometric distributions on the subset selection, and subsequently corrected these using the Benjamini-Hochberg FDR procedure, setting the FDR-value threshold to 0.1. Both over-representation and under-representation were accounted for in the analyses. The choice of background, i.e., the full set against which the subset is compared, is crucial for obtaining accurate results. For this, we used all mapped and processed proteins with well-defined *t*-tests based on the three biological replicates.

We collected GO term information for *Pseudomonas aeruginosa* using UniProt, release: 2024_03 (UniProt Consortium, 2023), and mapped the GO term IDs onto the Gene Ontology Knowledgebase using AmiGO 2, version: 2.5.17 (Carbon and Mungall, 2024), aligning the information with the UniProt IDs. Out of 3811 proteins, 3253 were associated with GO terms, and the remaining were assigned a placeholder value to ensure their inclusion in the analyses, since the non-annotated proteins may be of particular interest for further study.

In the results, we visualized the enriched GO terms using volcano plots. The x-coordinate of each GO term represents the log2 value of its enrichment (blue indicates under-representation and red indicates over-representation), and the range is fixed and visually capped at +/- 2.5. The y-coordinate represents the FDR values associated with each GO term (not to be confused with the protein-level FDR-values). The area of the GO-term circles is proportional to the number of proteins associated with the GO term in the subset. All terms with a GO-term FDR value > 0.1 are indicated in grey and are excluded from further analysis.

### 2.7 Resources and tools

Data analyses and visualizations were performed using Python 3.11, along with standard modules such as numpy, pandas, scipy, matplotlib, and seaborn, among others. The *t*-tests and dose-correlation statistics were conducted using R.

## 3 Results

Here, we present the results of our comparative proteomic investigation between the *Pseudomonas aeruginosa* world epidemic clinical strain and the wildtype control strain under four increasing concentrations (in mg/L): 1.6, 4, 8, 16 and 0.2, 0.5, 1.0, 2.0, which equates to 0.1x, 0.25x, 0.5x, and 1.0x MIC for the clinical and control strains respectively. The proteome data, acquired and processed as described in the Materials and Methods section, comprised a total of *N*=3811 proteins. For each protein, we calculated the LFQ intensity values across the five concentrations for both strains. Subsequently, a *t*-test was performed across the biological replicates, using the control probe (no antibiotic) experiments as controls, to derive protein-specific *p*-values.

### 3.1 Results overview

Exploration of the overall trends in the data, as visualized in **Figure 2**, suggests that the protein abundance in the clinical strain remains stable under the influence of imipenem, unlike the control strain. The PCA plot (**Figure 2a**) reveals that the largest variance between the experiments is attributed to the strains themselves (PC1, x-axis). The second largest factor of variance (PC2, y-axis) corresponds to the increasing imipenem concentration, as evidenced by the gradual transition from control probe (no antibiotic) in the top-right to 1.0x MIC in the bottom-right for the control strain. The heatmap (**Figure 2b**) displays a highly intercorrelated block in the top-left corner, representing the clinical strain experiments. The bottom-right corner shows a slight but clear gradient towards covariance among experiments with similar imipenem concentrations. Notably, there is high similarity between the strains, with correlation coefficients around 0.85 for the control probes (A1 and B1) and 0.80 at 1.0x MIC (A5 and B5).

### 3.2 Proteomic changes within strains

Based on the overall results from **Figure 2**, we expect the significant fold changes for individual proteins across MIC concentrations to be more pronounced for the control strain than the clinical strain. The volcano plots in **Figure 3** confirms this prospect. Here, we have plotted the *N*=3811 proteins with a fold change threshold of 1.5 for both the control (**Figure 3a**) and clinical (**Figure 3b**) strains at 1.0x MIC, as compared with a *t*-test to 0.0x MIC (see Materials and Methods for details).

**Figure 3:**
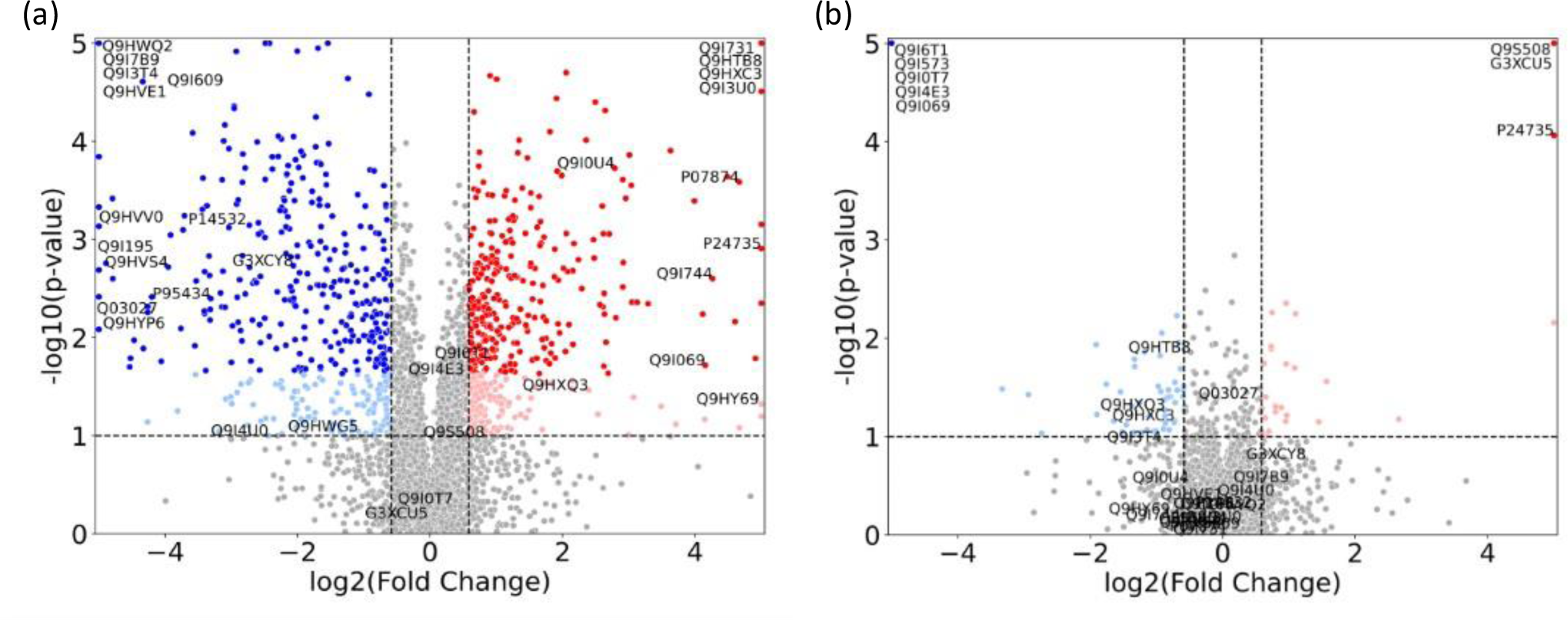
Volcano plots illustrating the proteomic changes in Pseudomonas aeruginosa for 1.0x MIC relative to control probe (no imipenem) for (a) the control strain (ATCC 27853) and (b) the clinical strain (ST235) with p-values along the y-axis and log2 fold change along the x-axis. Proteins with a fold change < 1.5 in either direction, or a p-value > 0.1 are depicted in grey. Light colors represent proteins with p-values < 0.1, but with FDR corrected p-values > 0.1, while full colors represent FDR corrected p-values < 0.1. Significant upregulated proteins are shown in red, while relevant downregulated proteins are shown in blue. Specific proteins of interest are indicated by their UniProt ID and constitute the union of the proteins in **Table 2** and **Table 6**.

**Table 2:**
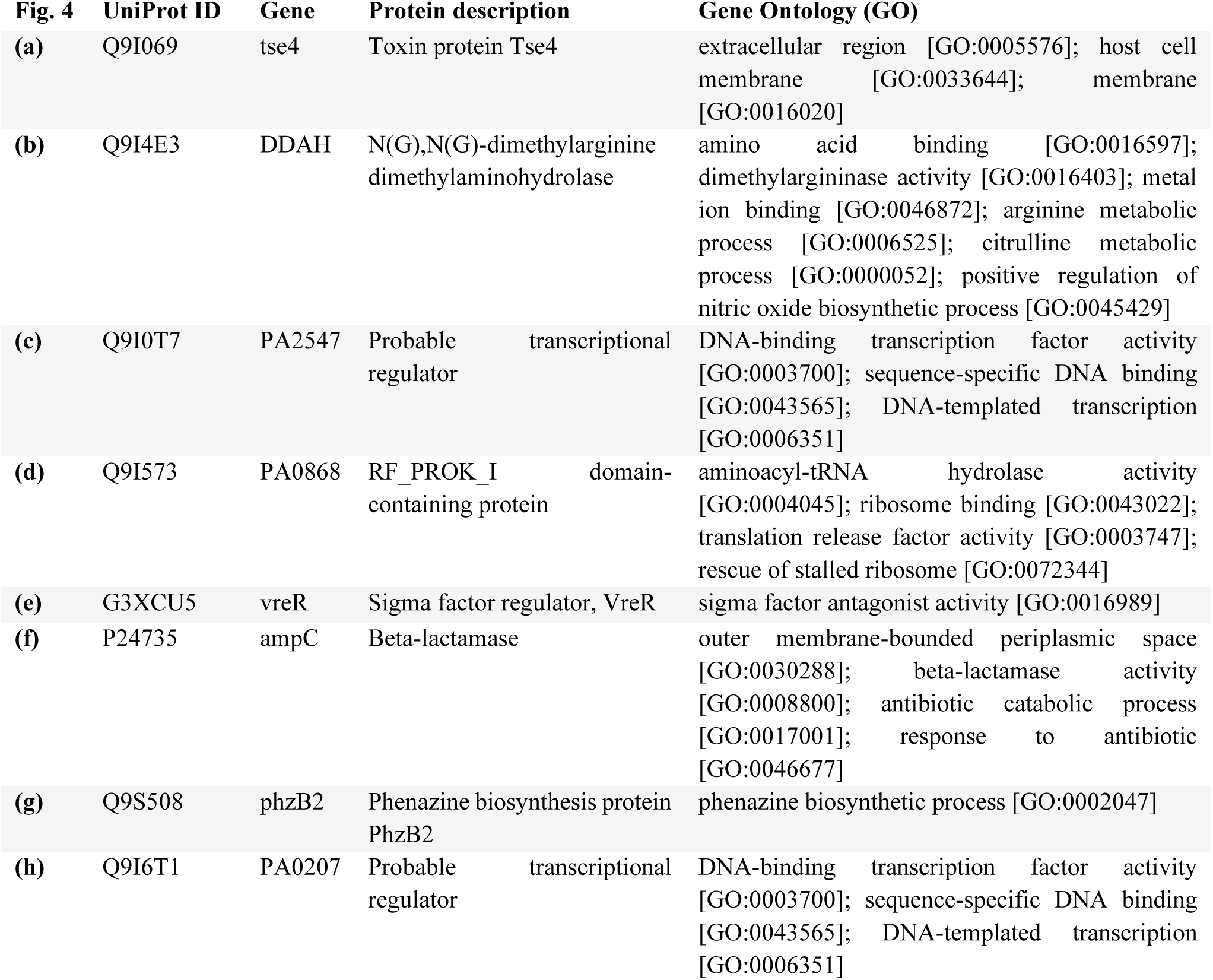
Significant proteins in the clinical strain with associated GO terms.

In these plots, an increase in fold change from 0.0x MIC to 1.0x MIC, i.e. up-regulation, is depicted by red dots, while a decrease in fold change, i.e. down-regulation, is represented by blue dots. Each dot corresponds to a protein. For both strains, we display the proteins with *p*-values above 0.1, and fold change below the threshold, as grey, while light red and blue denote *p*-values below 0.1 and full (darker) red and blue depict the proteins with FDR corrected *p*-values below 0.1. As expected, the control strain shows a range of significant FDR values, while the clinical strain only has 8 proteins fulfilling the criteria: AmpC (P24735), PA0868 (Q9I573), DDAH (Q9I4E3), Tse4 (Q9I069), PA2547 (Q9I0T7), PA0207 (Q9I6T1), VreR (G3XCU5) and PhzB2 (Q9S508). These proteins (depicted in the top corners of **Figure 3b**) are further explored in **Figure 4** and **Table 2** and detailed results will be presented in the following subsection.

**Figure 4:**
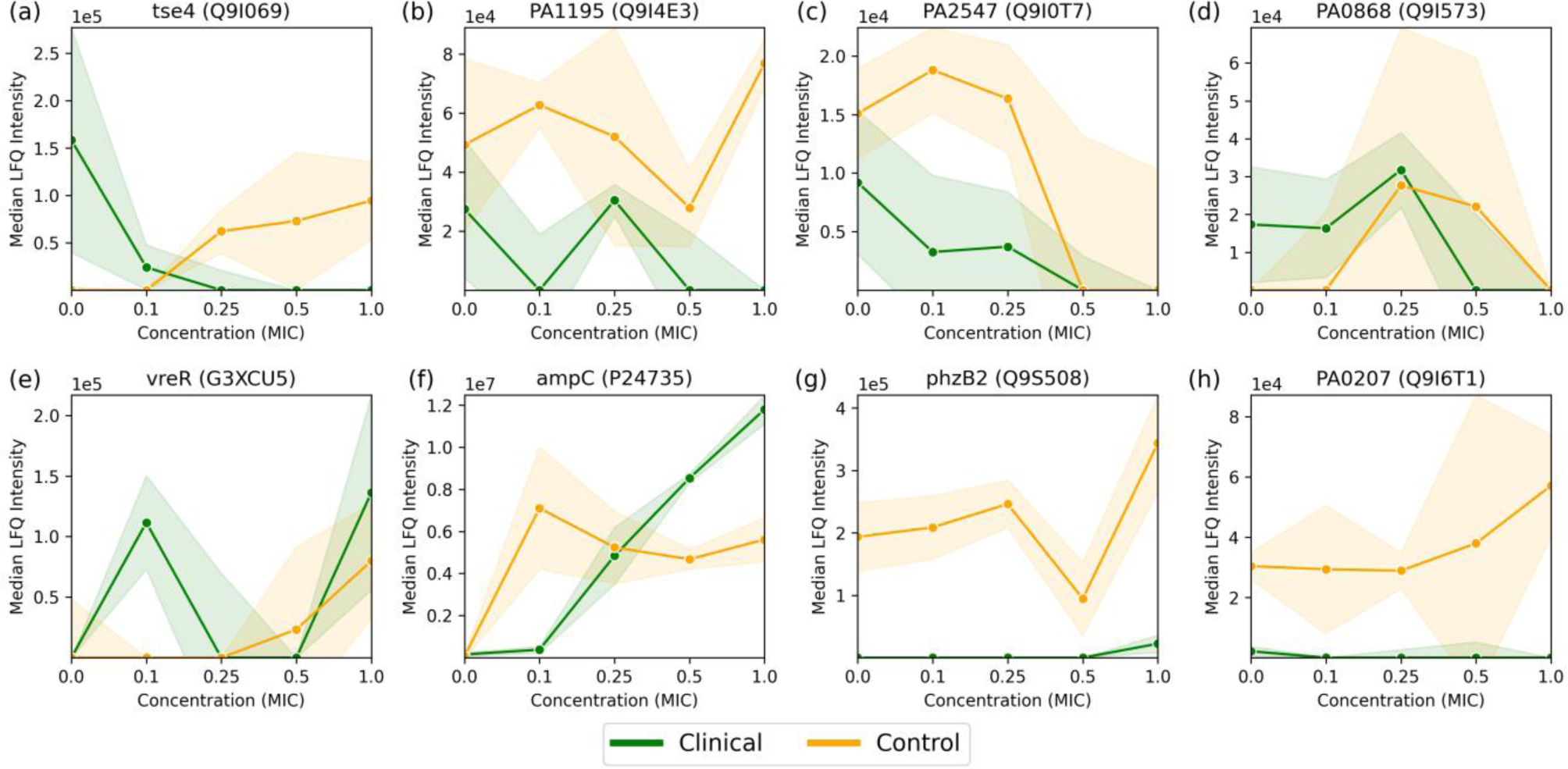
Median LFQ intensity scores plotted against increasing fractions of MIC, ranging from no antibiotic (zero concentration) to 1.0x MIC, for 8 proteins (a-h) passing the FDR corrected t-test for the Pseudomonas aeruginosa clinical strain, as displayed in **Figure 3b**. The clinical strain is shown in green, the control strain is shown in orange, and the shaded area depicts the associated standard deviations (n=3) among the replicates with a linear interpolation between the measurements.

For the normalized 1.0x MIC control strain experiments, displayed in **Figure 3a**, there are *n*=601 proteins with FDR value below 0.1 which show a fold change greater than 1.5 (depicted in dark red and blue), and *n*=1219 proteins adhere to the fold change threshold alone. The respective numbers for the clinical strain are *n*=8 (where *n*=79 have an uncorrected *p*-value below 0.1) and *n*=358, as visualized in **Figure 3b**.

### 3.3 Comparison of specific proteins

As mentioned in the previous section, 8 proteins passed the FDR value threshold of 0.1 for the clinical strain, making them resistome candidates which are significantly involved in the antimicrobial resistance in *Pseudomonas aeruginosa*. These proteins are shown in **Figure 4** and **Table 2**, where **Figure 4a-d** show an overall down-regulation of the specific protein in the clinical strain (in green) for increasing MIC, **Figure 4e-f** show an overall up-regulation for increasing MIC, while **Figure 4g-h** show a particularly large difference between the strains, i.e., where the LFQ intensity in the clinical strain is much lower than the control strain. The control strain (in orange) shows an overall trend towards up-regulation for these 8 proteins, aside from the proteins in **Figure 4c** and **4d**. The standard deviations, indicated by shaded green for the clinical strain and shaded orange for the control, vary. Large standard deviations, such as for PA0868 (Q9I573) in **Figure 4d**, indicate high variability in protein expression, while the high-LFQ intensity protein AmpC (P24735) in **Figure 4f** displays a clear signal, suggesting consistent protein expression across replicates. The dose-correlation statistics were calculated for all proteins for both strains, but since the number of measurements per protein is low, the associated *p*-values are poor, potentially making the correlation values misleading. Consequently, the specific dose correlations values will not be discussed here, but please refer to Supplementary Tables S1 and S2 for details on clinical and control strains respectively.

The proteins in **Figure 4** are described in more detail in **Table 2**, where a description and associated gene ontologies are added (see Materials and Methods for details). These findings and their implications for understanding antimicrobial resistance in *Pseudomonas aeruginosa* will be discussed further in the Discussion section.

### 3.4 GO term enrichment within strains

To better understand the biological implications of the significant up-and downregulated protein subsets (highlighted in red and blue in **Figure 3**), we conducted a GO term enrichment analysis as outlined in the Materials and Methods section. This analysis allows us to assess the biological relevance of our protein subsets, indicated by the enrichment of biological traits, when compared to the original full set of *N*=3811 proteins. By comparing high-fold-change proteins between 1.0x MIC and control probe (no imipenem), we can identify the biological functions and processes that the organism upregulates or downregulates. This comparison also provides *p*-values, which can furthermore be FDR corrected, indicating the statistical significance of these enrichments. I.e., there are two levels of (corrected) *p*-values at this stage: (i) protein-level *p*-values, based on *t*-tests for individual proteins, as depicted in **Figure 3**, and (ii) protein-subset level *p*-values, where enrichments of protein annotations in the subset compared to the full, original set are assessed via hypergeometric processes. Thus, a random selection of proteins, which does not reflect real biology, is expected to meet the protein-subset level, GO-term FDR threshold of 0.1, thus reducing the overall likelihood of false positive GO-term enrichment scores at 10%. In the following, we present results based on the union of upregulated (red) and downregulated (blue) proteins, as depicted in **Figure 3**. GO term analysis was performed on up-and downregulated proteins separately, but these results were too similar to the union set, when combined, to warrant a separate discussion.

The GO term enrichment analyses were conducted for both the control and clinical strains of *Pseudomonas aeruginosa*, as depicted in **Figure 5**. The protein subsets in **Figure 5a-d** were all subject to a fold change threshold of 1.5 per protein. Furthermore, **Figure 5a** has a protein-level FDR-value threshold of 0.1 applied to it, and **Figure 5b** an uncorrected protein-level *p*-value threshold of 0.1. The reason for this choice is based on numbers, since the clinical strain only had 8 proteins adhering to the FDR threshold, which is deemed too small for a sensible GO term analysis for comparative purposes. **Figures 5c-d** has no additional *p*-value thresholds and is only subject to a fold-change threshold of 1.5. The protein subsets in **Figures 5a** and **5b** are thus based on the *n*=601 and *n*=79 proteins, corresponding to the dark red and dark blue dots in **Figure 3a**, and the light and dark red and blue dots in **Figure 3b** respectively. In **Figures 5c**, and **5d**, on the other hand, the protein subsets comprise *n*=1219 and *n*=358 proteins respectively, equating to all the dots outside the dashed vertical fold-change threshold lines. The significantly enriched GO terms, with hypergeometric subset enrichment FDR values < 0.1, are indicated by circles color-coded as red for over-represented terms compared to what you would expect from random processes (enrichment above 1) and blue for under-represented ones (enrichment below 1). The size of the circles indicates the number of proteins associated with each GO term, and the terms are alphabetically labeled based on their significance (decreasing FDR values).

**Figure 5:**
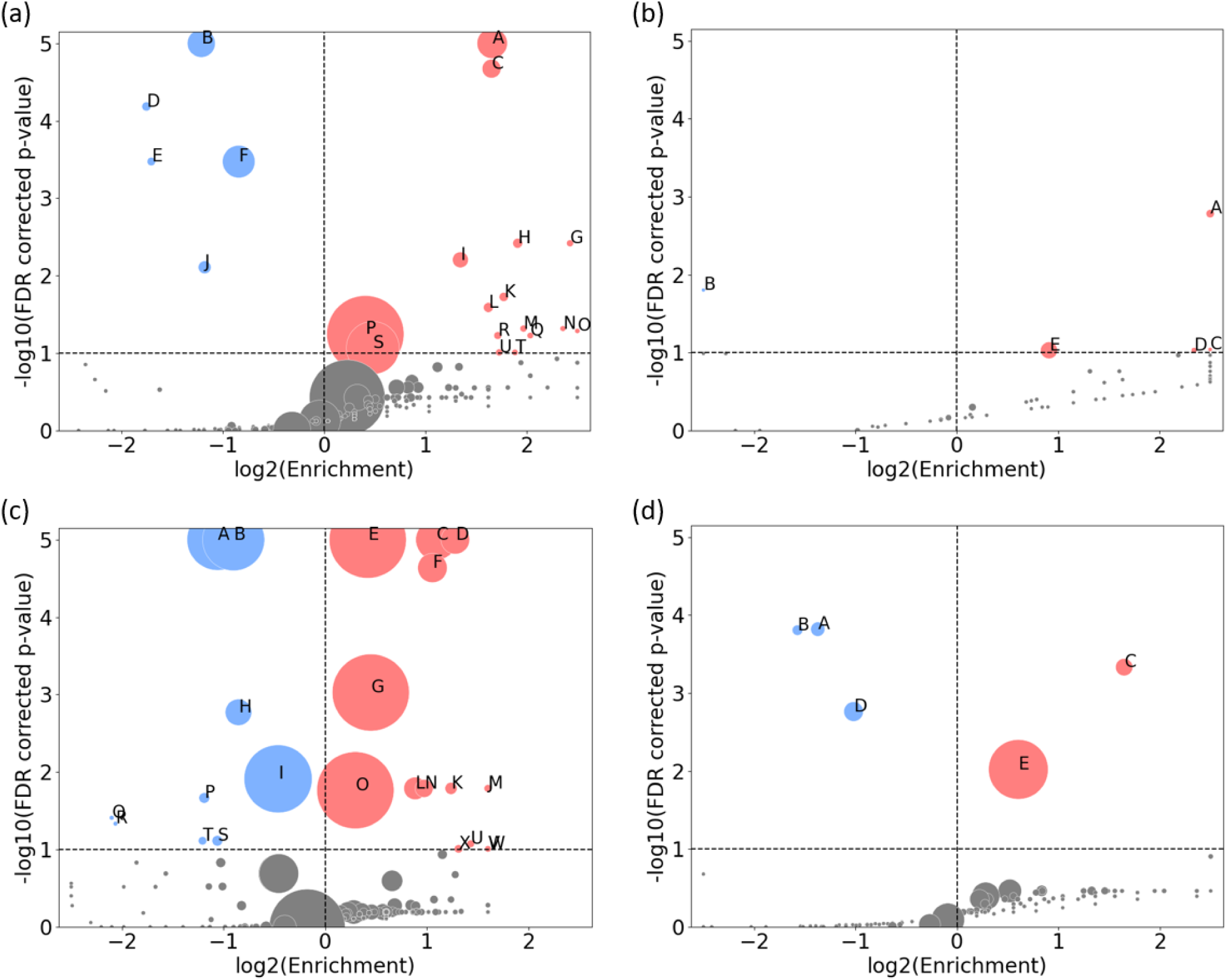
Comparative GO term enrichment in Pseudomonas aeruginosa strains. The volcano plots depict the enrichment of GO terms in protein subsets from the control strain (a, c) and the clinical strain (b, d), with: (a) a protein-level FDR threshold of 0.1, (b) a protein-level p-value threshold of 0.1, and (c, d) no p-value threshold. All subsets adhere to a protein-level fold-change threshold of 1.5. The x-axis represents the log2 enrichment score, while the y-axis shows the transformed FDR. GO terms are represented by circles, where the size corresponds to the number of proteins linked to each GO term, visualizing the difference of significant proteins associated with the control strain (left) compared to the clinical strain (right). The circles are color-coded as light red for over-represented (enriched) and light blue for under-represented (“under-enriched”) terms. **Table 3** and **Table 4** provide additional information on the terms labeled in capital letters for (a) and (d) respectively.

**Table 3:** GO terms enriched in the control strain for protein *t*-test FDR < 0.1 and fold change > 1.5 (label from Figure 5a)

| Label | GO ID | Enrichment | Size | FDR | GO term |
| --- | --- | --- | --- | --- | --- |
| ● A | GO:0042597 | 3.158 | 38 | 1.39e-09 | periplasmic space |
| ● B | GO:0005829 | 0.431 | 35 | 3.57e-08 | cytosol |
| ● C | GO:0030288 | 3.143 | 23 | 2.12e-05 | outer membrane-bounded periplasmic space |
| ● D | GO:0003700 | 0.296 | 10 | 6.55e-05 | DNA-binding transcription factor activity |
| ● E | GO:0003677 | 0.306 | 9 | 3.37e-04 | DNA binding |
| ● F | GO:0005737 | 0.557 | 41 | 3.37e-04 | cytoplasm |
| ● G | GO:1904680 | 5.381 | 7 | 3.82e-03 | peptide transmembrane transporter activity |
| ● H | GO:0002049 | 3.758 | 11 | 3.82e-03 | pyoverdine biosynthetic process |
| ● I | GO:0043190 | 2.54 | 19 | 6.25e-03 | ATP-binding cassette (ABC) transporter complex |
| ● J | GO:0006355 | 0.441 | 15 | 7.81e-03 | regulation of DNA-templated transcription |
| ● K | GO:0038023 | 3.417 | 10 | 0.0189 | signaling receptor activity |
| ● L | GO:0015562 | 3.075 | 11 | 0.0258 | efflux transmembrane transporter activity |
| ● M | GO:0006826 | 3.913 | 7 | 0.0483 | iron ion transport |
| ● N | GO:0042938 | 5.125 | 5 | 0.0483 | dipeptide transport |
| ● O | GO:0015419 | 6.15 | 4 | 0.0518 | ABC-type sulfate transporter activity |
| ● P | GO:0005886 | 1.325 | 100 | 0.0568 | plasma membrane |
| ● Q | GO:0015891 | 4.1 | 6 | 0.0594 | siderophore transport |
| ● R | GO:0005615 | 3.28 | 8 | 0.0594 | extracellular space |
| ● S | GO:0016020 | 1.394 | 68 | 0.0856 | membrane |
| ● T | GO:0033103 | 3.69 | 6 | 0.0982 | protein secretion by the type VI secretion system |
| ● U | GO:1990281 | 3.311 | 7 | 0.0982 | efflux pump complex |

**Table 4:** GO terms enriched in the clinical strain with fold change > 1.5 (label from Figure 5d)

| Label | GO ID | Enrichment | Size | FDR | GO term |
| --- | --- | --- | --- | --- | --- |
| ● A | GO:0005737 | 0.386 | 17 | 1.51e-04 | cytoplasm |
| ● B | GO:0005524 | 0.336 | 12 | 1.56e-04 | ATP binding |
| ● C | GO:0042597 | 3.131 | 21 | 4.65e-04 | periplasmic space |
| ● D | GO:0005829 | 0.493 | 24 | 1.73e-03 | cytosol |
| ● E | NOGO | 1.52 | 77 | 9.55e-03 | No known GO terms |

**Table 3** provides a list of the significantly enriched GO terms shown in **Figure 5a** with their respective statistics. Noteworthy terms include proteins associated with various cellular components (e.g., periplasmic space, cytosol, cytoplasm, and membrane), transporter activities (peptide transmembrane, efflux transmembrane, ABC-type sulfate and siderophore), DNA interactions, and specific processes (e.g., pyoverdine biosynthesis and iron ion transport). The clinical strain where only the fold-change threshold is applied, displayed in **Figure 5d** and **Table 4**, exhibits some of the same terms as in **Table 3**, albeit with lesser significance. A unique feature here is the enrichment of proteins without associated GO term annotations, which, with an enrichment score of 1.52, is still significant due to the substantial number of proteins in the “NOGO” pseudo-term as GO annotations are missing.

The corresponding results for **Figures 5b** and **5c** are presented in Supplementary Tables S3 and S4 respectively. The apparent overlap between these results and those listed in Tables 4 and 5 indicates that our approach is robust and reliable, even when using different or no *p*-value thresholds.

**Table 5:** GO terms enriched in differential protein expression (label from Figure 6d)

| Label | GO ID | Enrichment | Size | FDR | GO term |
| --- | --- | --- | --- | --- | --- |
| ● **A** | GO:0042597 | 6.158 | 18 | 4.02e-08 | periplasmic space |
| ● **B** | GO:0005829 | 0.152 | 3 | 1.92e-04 | cytosol |
| ● **C** | GO:0031460 | 10.126 | 4 | 0.0155 | glycine betaine transport |
| ● **D** | GO:0016989 | 10.126 | 4 | 0.0155 | sigma factor antagonist activity |
| ● **E** | GO:0043190 | 4.403 | 8 | 0.0155 | ATP-binding cassette (ABC) transporter complex |
| ● **F** | GO:0042121 | 8.438 | 5 | 0.0155 | alginic acid biosynthetic process |
| ● **G** | GO:0071475 | 12.658 | 3 | 0.0263 | cellular hyperosmotic salinity response |
| ● **H** | GO:0015838 | 12.658 | 3 | 0.0263 | amino-acid betaine transport |
| ● **I** | GO:0015226 | 12.658 | 3 | 0.0263 | carnitine transmembrane transporter activity |
| ● **J** | GO:1902495 | 25.315 | 2 | 0.0295 | transmembrane transporter complex |
| ● **K** | GO:0010438 | 25.315 | 2 | 0.0295 | cellular response to sulfur starvation |
| ● **L** | GO:0005576 | 4.467 | 6 | 0.0319 | extracellular region |
| ● **M** | NOGO | 1.622 | 33 | 0.0405 | No known GO terms |
| ● **N** | GO:0030254 | 6.329 | 4 | 0.0429 | protein secretion by the type III secretion system |
| ● **O** | GO:0051131 | 16.877 | 2 | 0.0574 | chaperone-mediated protein complex assembly |
| ● **P** | GO:0015418 | 16.877 | 2 | 0.0574 | ABC-type quaternary ammonium compound transporting activity |
| ● **Q** | GO:0004130 | 12.658 | 2 | 0.0986 | cytochrome-c peroxidase activity |
| ● **R** | GO:0033644 | 12.658 | 2 | 0.0986 | host cell membrane |

### 3.5 Differential protein expression and GO term enrichment

To understand the proteomic landscape of imipenem resistance in *Pseudomonas aeruginosa*, we have thus far delved into within-strain comparisons, specifically between the 1.0x MIC and control probe (no antibiotic) experiments. We have also cross-compared these results between the clinical and control strains. Now, we consider a more direct comparison across strains, by using the difference between the normalized, log2-transformed LFQ values of the control and clinical strains as a measure of fold-change (see Materials and Methods for details).

The volcano plots depicted in **Figures 6a** and **6c** illustrate fold-change thresholds of 1.5 and 8 respectively (equivalent to approximately +/- 0.585 and exactly +/- 3 in log2 space respectively). The reason for these choices stems from **Figure 3**, where we wanted a comparable example (**Figure 6a**) as well as a suitably sparse counter example, where only proteins with large differences in expression are included (**Figure 6c**). Other thresholds could be chosen, but we found a >8-fold change threshold to be a good compromise between sparsity and representativeness. The FDR corrected *p*-values displayed on the y-axis are derived from the control strain *t*-test. These are included for visual comparability with **Figure 3** and are not considered in the GO term enrichment analysis displayed in **Figure 6b** and **6c**.

**Figure 6:**
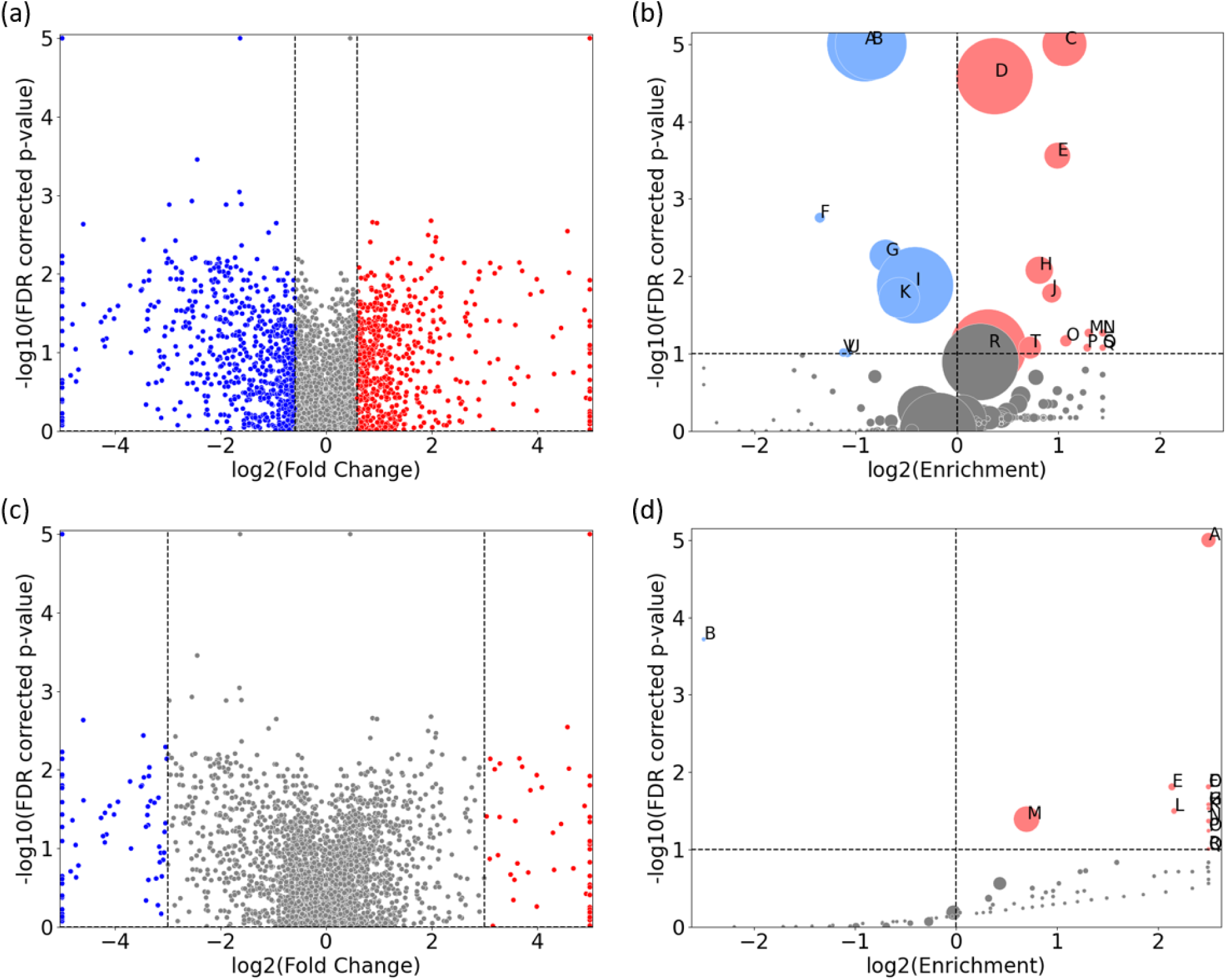
Direct comparison of protein expression between the control-probe-normalized control strain and clinical strain for 1.0x MIC imipenem treatment. (a) and (c) are volcano plots displaying the normalized differential protein intensity fold-change along the x-axis, with a fold-change threshold of 1.5 and 8, respectively. Red dots have larger positive fold change in the control strain, from 1.0x MIC relative to control probe (no antibiotic), compared to the clinical strain, and, correspondingly, larger negative fold change for the blue dots. (b) and (d) are GO term volcano plots for the protein subset based on the non-grey proteins in (a) and (b) respectively. The FDR values along the y-axis in (a) and (c) are derived from the control strain and applied for visual comparison.

As **Figure 2** demonstrates, the clinical strain’s overall protein expression is not significantly affected by increasing MIC. Consequently, we anticipate that the normalized differential protein expression will mirror the results observed in the control strain. This expectation is confirmed in **Figure 6b**, which bears both visual and content similarities to **Figure 5c** (detailed information can be found in Supplementary Table S5). A noteworthy change, however, is the enrichment of proteins associated with magnesium ion binding.

A GO term enrichment analysis was conducted using the 146 proteins with fold change of more than 8, denoted by blue and red dots in **Figure 6c**. The results of this analysis are presented in **Figure 6c** and further detailed in **Table 5**. The proteins involved are provided in Supplementary Table S6 (in the “IDs” column).

Several of these terms are recognizable from protein subsets seen in e.g. **Table 3**, but there are also differences from earlier protein subsets. These include transport terms like glycine betaine, amino-acid betaine, and carnitine transmembrane transport. There are specific enzyme-related terms such as sigma factor antagonist and cytochrome-c peroxidase, along with chaperone-mediated protein complex assembly. The list also includes cellular responses to hyperosmotic salinity and sulfur starvation, protein secretion by the type III secretion system, alginic acid biosynthesis, and structures like the extracellular region and host cell membrane. It is also important to highlight the *n*=33 proteins not associated with any known GO terms, representing a significant over-representation. These proteins, along with other specific proteins, will be the focus of our investigation in the subsequent section.

### 3.6 Comparison of selected proteins with large differential protein expression between strains

In **Figure 6**, we visualized the differential protein expression comparing the difference in the clinical strain to that of the control strain, and in **Figure 6c**, we applied a threshold demanding this difference to be >8-fold. Here, we will delve into some of the proteins adhering to this threshold. To obtain a manageable number of proteins, we employed some criteria for selection: (i) low or negative correlation values (< 0.4) between clinical and control strain across the control probe (no antibiotic) plus 4 imipenem concentrations (0.1x, 0.25x, 0.5x and 1.0x MIC), (ii) striking differences in expression, and/or (iii) part of significantly enriched GO terms in **Table 5**. This process led us to 26 proteins, 24 of which are shown in **Figure 7**. The remaining two proteins are Tse4 (Q9I069), already displayed in **Figure 4a**, and PpiC2 (Q9HWK5) which is highly expressed in the clinical strain, and comparably non-expressed in the control across all concentrations. The remaining 24 proteins are split so that **Figure 7a-o** has an overall downregulation for the control for increasing imipenem concentrations, while **Figure 7p-x** show an overall up-regulation for the control. The clinical strain displays more stability across varying concentrations, which is not surprising given the results shown in **Figure 2**. However, **Figures 7s** and **7w**, alongside **Figure 4a**, displaying the proteins Wzm (Q9HTB8), PA0101 (Q9I731) and Tse4 (Q9I069) respectively, also show clear down-regulation in the clinical strain for increasing concentrations, in addition to the up-regulation in the control strain. As in **Figure 4**, the standard deviations are shown as green (clinical) and orange (control) shading, where linear interpolation has been used between the measurements. While several proteins display large standard deviations for the clinical strain, most control strain trends are associated with comparably small standard deviations. This indicates a clear biological effect of increasing concentrations of imipenem for the *Pseudomonas aeruginosa* (ATCC 27853) control strain which can be related to these proteins and their associated functions and compartments. More information about the proteins in **Figure 7** can be found in **Table 6**, including their UniProt description (UniProt Consortium, 2023) and associated GO terms (Carbon and Mungall, 2024). Some proteins, PA2384 (Q9I195), PA4129 (Q9HWQ2), PA1469 (Q9HVV0), PA1034 (Q9I4U0), PA1404 (Q9I3U0) and PA0101 (Q9I731), in **Figure 7c, d, i, m, p** and **w** respectively, are uncharacterized and have thus no associated GO terms. These results will be further analyzed in the Discussion section, where we will shed light on their potential association with imipenem and antimicrobial activity.

**Figure 7:**
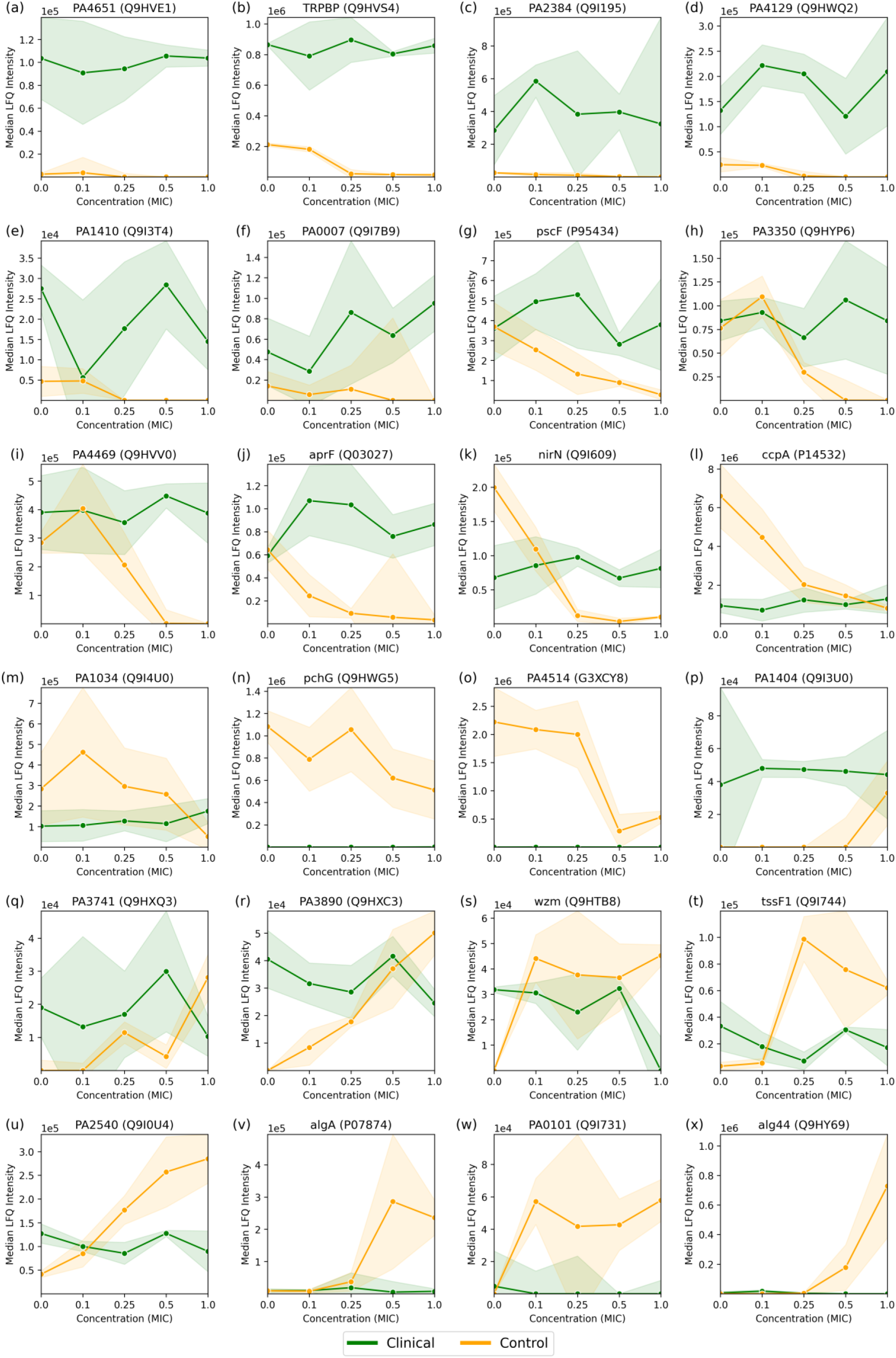
Median LFQ intensity scores plotted against increasing fractions of the MIC, ranging from no imipenem (zero concentration) to 1.0x MIC, for 24 proteins (a-x) with particularly large and/or interesting differential protein expressions between the Pseudomonas aeruginosa clinical and control strains, as displayed by red and blue dots in **Figure 6c**. The clinical strain is shown in green, the control strain is shown in orange, and the shaded area depicts the associated standard deviations (n=3) among the replicates with a linear interpolation between the measurements.

**Table 6:** Selected proteins with >8-fold difference between strains at 1.0x MIC.

| Fig. 7 | UniProt ID | Gene | Protein description | Gene Ontology (GO) |
| --- | --- | --- | --- | --- |
| (a) | Q9HVE1 | PA4651 | Probable pili assembly chaperone | outer membrane-bounded periplasmic space [GO:0030288]; cell wall organization [GO:0071555] |
| (b) | Q9HVS4 | TRPBP | Tripeptide-binding protein | ATP-binding cassette (ABC) transporter complex [GO:0043190]; outer membrane-bounded periplasmic space [GO:0030288]; peptide transmembrane transporter activity [GO:1904680]; dipeptide transport [GO:0042938]; protein transport [GO:0015031] |
| (c) | Q9I195 | PA2384 | Uncharacterized protein |  |
| (d) | Q9HWQ2 | PA4129 | Uncharacterized protein |  |
| (e) | Q9I3T4 | PA1410 | Probable periplasmic spermidine/putrescine-binding protein | periplasmic space [GO:0042597]; polyamine binding [GO:0019808]; polyamine transport [GO:0015846] |
| (f) | Q9I7B9 | PA0007 | Cytochrome c domain-containing protein | cytochrome-c peroxidase activity [GO:0004130]; electron transfer activity [GO:0009055]; heme binding [GO:0020037]; metal ion binding [GO:0046872] |
| (g) | P95434 | pscF | Type III needle protein PscF | cell surface [GO:0009986]; extracellular region [GO:0005576]; type III protein secretion system complex [GO:0030257]; protein secretion by the type III secretion system [GO:0030254] |
| (h) | Q9HYP6 | PA3350 | Flagella basal body P-ring formation protein FlgA | periplasmic space [GO:0042597]; bacterial-type flagellum assembly [GO:0044780] |
| (i) | Q9HVV0 | PA4469 | Uncharacterized protein |  |
| (j) | Q03027 | aprF | Alkaline protease secretion protein AprF | cell outer membrane [GO:0009279]; efflux pump complex [GO:1990281]; efflux transmembrane transporter activity [GO:0015562]; porin activity [GO:0015288]; secretion [GO:0046903] |
| (k) | Q9I609 | nirN | Dihydro-heme d1 dehydrogenase | periplasmic space [GO:0042597]; electron transfer activity [GO:0009055]; heme binding [GO:0020037]; metal ion binding [GO:0046872]; oxidoreductase activity [GO:0016491]; heme biosynthetic process [GO:0006783] |
| (l) | P14532 | ccpA | Cytochrome c551 peroxidase | periplasmic space [GO:0042597]; cytochrome-c peroxidase activity [GO:0004130]; electron transfer activity [GO:0009055]; heme binding [GO:0020037]; metal ion binding [GO:0046872] |
| (m) | Q9I4U0 | PA1034 | Uncharacterized protein |  |
| (n) | Q9HWG5 | pchG | Pyochelin biosynthetic protein PchG | nucleotide binding [GO:0000166] |
| (o) | G3XCY8 | PA4514 | Probable outer membrane receptor for iron transport | cell outer membrane [GO:0009279]; pore complex [GO:0046930]; porin activity [GO:0015288]; siderophore uptake transmembrane transporter activity [GO:0015344]; signaling receptor activity [GO:0038023]; iron ion transport [GO:0006826]; siderophore transport [GO:0015891] |
| (p) | Q9I3U0 | PA1404 | Uncharacterized protein |  |
| (q) | Q9HXQ3 | PA3741 | HotDog ACOT-type domain-containing protein | cytoplasm [GO:0005737]; cytosol [GO:0005829]; long-chain fatty acyl-CoA binding [GO:0036042]; long-chain fatty acyl-CoA hydrolase activity [GO:0052816]; acyl-CoA metabolic process [GO:0006637]; fatty acid catabolic process [GO:0009062] |
| (r) | Q9HXC3 | PA3890 | Probable permease of ABC transporter | ATP-binding cassette (ABC) transporter complex [GO:0043190]; transmembrane transporter complex [GO:1902495]; carnitine transmembrane transporter activity [GO:0015226]; amino-acid betaine transport [GO:0015838]; cellular hyperosmotic salinity response [GO:0071475]; glycine betaine transport [GO:0031460] |
| (s) | Q9HTB8 | wzm | Transport permease protein | ATP-binding cassette (ABC) transporter complex [GO:0043190]; inner membrane pellicle complex [GO:0070258]; ABC-type transporter activity [GO:0140359]; lipopolysaccharide transport [GO:0015920]; O antigen biosynthetic process [GO:0009243]; transmembrane transport [GO:0055085] |
| (t) | Q9I744 | tssF1 | Type VI secretion system component TssF1 |  |
| (u) | Q9I0U4 | PA2540 | Bifunctional alpha/beta hydrolase/class I SAM-dependent methyltransferase | membrane [GO:0016020]; lipase activity [GO:0016298] |
| (v) | P07874 | algA | Alginate biosynthesis protein AlgA | GTP binding [GO:0005525]; mannose-1-phosphate guanylyltransferase (GTP) activity [GO:0004475]; mannose-6-phosphate isomerase activity [GO:0004476]; alginic acid biosynthetic process [GO:0042121]; GDP-mannose biosynthetic process [GO:0009298] |
| (w) | Q9I731 | PA0101 | Uncharacterized protein |  |
| (x) | Q9HY69 | alg44 | Mannuronan synthase | Gram-negative-bacterium-type cell wall [GO:0009276]; periplasmic space [GO:0042597]; alginate synthase activity [GO:0047643]; cyclic-di-GMP binding [GO:0035438]; long-chain fatty acid-CoA ligase activity [GO:0004467]; alginic acid biosynthetic process [GO:0042121]; single-species biofilm formation [GO:0044010] |

## 4 Discussion

The proteomic landscape of *Pseudomonas aeruginosa* under the influence of imipenem, a commonly used carbapenem antibiotic, has been the subject of investigation in this study revealing key insights into the molecular mechanisms of antimicrobial resistance. Our comparative analysis between the world epidemic clinical strain and the control strain has highlighted changes in the proteome that occur in response to varying imipenem concentrations. Utilizing normalized label-free quantification (LFQ) intensities, protein group level *t*-tests, and Gene Ontology (GO) enrichment analysis of protein subsets, we have been able to delve deeper into the molecular mechanisms of imipenem resistance.

The observed differences in minimum inhibitory concentrations (MICs) between the clinical and control strains (**Table 1**; **Figure 2**) underscore the clinical strain’s enhanced resistance to imipenem, reflecting its ability to survive under antibiotic pressures that typically inhibit or kill susceptible strains (Leigue, Montiani-Ferreira and Moore, 2016). The varying antibiotic concentrations provide a suitable framework to explore the adaptive proteomic responses of both strains, revealing molecular adaptations associated with resistance.

In the clinical strain, eight proteins are identified as significant, mentioned in **Table 2**, each playing a role in the resistance strategy and with wide-reaching implications for understanding the mechanisms of imipenem resistance in *Pseudomonas aeruginosa*. The upregulation of beta-lactamase (AmpC, P24735) is consistent with its well-documented role in hydrolyzing beta-lactam antibiotics, including imipenem, and it is known to belong in a group of enzymes that provide multi-resistance to antibiotics (Jacoby, 2009). Its overexpression in the clinical strain could be a direct response to the antibiotic pressure, leading to the degradation of imipenem and thus, resistance (Berrazeg *et al*., 2015; Barbier *et al*., 2023; Elfadadny *et al*., 2024). In addition, also the sigma factor regulator VreR (G3XCU5), another upregulated protein, is likely involved in modulating the expression of genes that help the strain withstand antibiotic-induced stress, contributing to its resilience (Matilla *et al*., 2022).

Interestingly, Tse4 (Q9I069), a toxin protein associated with the Type VI secretion system, is downregulated in the clinical strain. This could potentially be caused by protein degradation. While RNA-seq data is not available for this study, future investigations comparing mRNA levels with protein expression could provide further insights. However, this could suggest a strategic conservation of resources, where the strain prioritizes essential survival functions over energy-intensive processes like toxin production under antibiotic stress. The associated GO terms, such as “beta-lactamase activity” (GO:0008800) and “sigma factor antagonist activity” (GO:0016989), highlight the biochemical pathways enhanced in response to imipenem, pointing to a multifaceted resistance mechanism involving both direct antibiotic degradation and stress response modulation. In contrast, the control strain exhibits a broad activation of cellular pathways in response to increasing imipenem concentrations.

Based on the Gene Ontology (GO) terms enriched in the *Pseudomonas aeruginosa* control strain comparing 1.0x MIC to the (no imipenem) control probes (**Figure 3a and Table 3**), there are several key processes of interest. The significant enrichment of GO terms related to membrane integrity and transport processes, such as “periplasmic space” (GO:0042597) and “efflux transmembrane transporter activity” (GO:0015562), suggests a defensive strategy focused on expelling the antibiotic and maintaining membrane stability (Young *et al*., 2019; Heywood and Lamont, 2020).

The upregulation of proteins involved in the “pyoverdine biosynthetic process” (GO:0002049) further indicates an enhancement of iron acquisition systems, potentially as a countermeasure to the oxidative stress imposed by imipenem. In addition, the production of pyoverdines, siderophores is essential for iron acquisition, growth and survival by mediating biofilm formation and pathogenicity (Dell’Anno *et al*., 2022). In addition to the broader activation of intracellular processes, as indicated by the enrichment of GO terms like “cytosol” (GO:0005829) and “cytoplasm” (GO:0005737), there is significant enrichment of terms related to the “ATP-Binding Cassette (ABC) Transporter Complex”, which may play a role as an antibiotic efflux transporter (Lewis, Ween and McDevitt, 2012). Also noteworthy is the involvement of Type VI Protein Secretion System Complex, often linked to biofilm production as an antagonistic mechanism (Hespanhol, Nóbrega-Silva and Bayer-Santos, 2023). Interestingly, some studies suggest that imipenem can inhibit biofilm production, which is associated with Type VI Secretion System (de Sousa *et al*., 2023). This further illustrates the control strain’s strategy to activate a broad spectrum of defensive pathways to cope with antibiotic threat.

As reflected by the results from **Table 4** (**Figure 5d**), the proteomic response in the clinical strain is more targeted, it involves fewer GO terms, reflecting stable resistance mechanism that does not require extensive proteasomal reconfigurations. The enrichment of “periplasmic space” (GO:0042597) and “cytosol” (GO:0005829) suggests that the clinical strain, like the control strain, emphasizes membrane integrity and cytoplasmic processes. However, the clinical strain does not show a broad spectrum of activated pathways, suggesting either pre-existing or well-adapted mechanisms that require little alteration in response to imipenem. What is remarkable, is the significant enrichment of uncharacterized proteins, indicated by the “NOGO” term, suggesting that the clinical strain may use novel or less-characterized proteins to its resistance strategy, e.g. by proteins like PA2384 (Q9I195) or PA4129 (Q9HWQ2) in **Figures 7c** and **7d**, to be further discussed below. This is indicative of potential areas where further research could uncover new aspects of antimicrobial resistance. If confirmed, and further investigated, the role of these proteins might underscore novel proteomic profiles linked to adaptation to antibiotic stress.

A direct comparison of protein expression between the clinical and control strains at 1.0x MIC reveal a clear difference in their strategies to cope with imipenem (**Table 5**, **Figure 6d**). The control strain shows substantial upregulation of proteins involved in membrane and transport processes, reflecting a reactive and broad-spectrum response to antibiotic pressure. The significant enrichment of GO terms like “glycine betaine transport” (GO:0031460) and “ABC transporter complex” (GO:0043190) suggests that the control strain activates mechanisms to manage osmotic stress and expel imipenem, respectively. Glycine betaine is known to protect cells against osmotic stress (Zhao *et al*., 2018), and its increased transport activity could be part of the control strain’s attempt to mitigate the effects of imipenem-induced stress. In contrast, the clinical strain shows enrichment in oxidative stress management pathways, such as “cytochrome-c peroxidase activity” (GO:0004130), highlighting its focus on specific oxidative stress management rather than broad reactive changes. The presence of “extracellular region” (GO:0005576) enrichment in the clinical strain suggests it may involve extracellular defense mechanisms, potentially modifying or interacting with its environment to resist the antibiotic. The pronounced differences in protein expression and overall cellular responses to imipenem between the clinical and control strains of *Pseudomonas aeruginosa* are further elucidated by the analysis of selected proteins with >8-fold differential expression, as detailed in **Table 6** and visualized in **Figure 7**. The clinical strain’s ability to maintain a relatively stable proteomic profile across varying concentrations of imipenem contrasts starkly with the control strain’s more reactive and fluctuating proteomic responses, which likely reflect a broad-spectrum attempt to mitigate antibiotic stress. Uncharacterized proteins, such as PA2384 (Q9I195) and PA4129 (Q9HWQ2), exhibit significant upregulation in the clinical strain, suggesting these proteins may be integral to its resistance mechanisms. The consistent expression of these proteins, even under high antibiotic pressure, indicates a potentially critical role in the clinical strain’s adaptation and survival. This upregulation, coupled with the stable expression of well-characterized proteins like AprF (Q03027) and AlgA (P07874), underscores the clinical strain’s reliance on pre-established resistance mechanisms, which seem to operate efficiently without the need for extensive proteomic reconfiguration in response to imipenem. In contrast, the control strain demonstrates significant proteomic shifts, particularly in proteins associated with membrane transport and stress response, such as PA4514 (G3XCY8) and Wzm (Q9HTB8). These shifts reflect the strain’s reactive adjustments to counteract the effects of imipenem, emphasizing a less specialized and more variable approach to resistance. The marked increase in expression of alginate biosynthesis proteins in the control strain at higher MICs also suggests an attempt to enhance biofilm formation as a defensive measure, a strategy that the clinical strain does not appear to rely on as heavily, likely due to its already optimized biofilm-related defenses.

These observations highlight the distinct proteomic strategies employed by the clinical and control strains. The clinical strain’s more stable and efficient resistance mechanisms, which involve both characterized and novel proteins, contrast with the control strain’s broader and more reactive defense responses. These findings suggest that the uncharacterized proteins in the clinical strain could be targets for future research aimed at uncovering novel antimicrobial resistance mechanisms and developing targeted therapies.

## 5 Conclusion

This study provides a comprehensive comparative analysis in *Pseudomonas aeruginosa* of the proteomic responses of a clinical strain, representative of the world epidemic clone (*P. aeruginosa* ST235), and a control strain (*P. aeruginosa* ATCC 27853) to varying concentrations of imipenem. The clinical strain’s ability to maintain a stable proteomic profile, even under high antibiotic pressure, underscores its reliance on pre-established resistance mechanisms. Both well-characterized and uncharacterized proteins have been found to be involved in the clinical strain’s adaptation and survival. This stability contrasts sharply with the control strain’s more reactive and fluctuating proteomic responses, which reflect a broad attempt to mitigate antibiotic stress. The control strain’s significant proteomic shifts, particularly in proteins associated with membrane transport and stress response, highlight its less specialized and more variable approach to resistance. These findings highlight the distinct proteomic strategies employed by the clinical and control strains. The clinical strain’s efficient and stable resistance mechanisms, involving both characterized and novel proteins, contrast with the control strain’s broader and more reactive defense responses. The identification of uncharacterized proteins in the clinical strain as potential resistome targets for future research underscores the importance of further investigation into novel antimicrobial resistance mechanisms. This could lead to the development of targeted therapies, providing new avenues in the fight against antimicrobial resistance.

## Supporting information

Supplementary Table S1: Dose-correlation statistics for control strain.

Supplementary Table S2: Dose-correlation statistics for clinical strain.

Supplementary Table S3: Tabular GO-term enrichment results corresponding to Figure 5b.

Supplementary Table S4: Tabular GO-term enrichment results corresponding to Figure 5c.

Supplementary Table S5: Tabular GO-term enrichment results corresponding to Figure 6b.

Supplementary Table S6: List of protein IDs associated with GO terms in Table 5.

## Declaration of generative AI and AI-assisted technologies in the writing process

During the preparation of this work the authors used ChatGPT-4 (via Microsoft Copilot) for proofreading and suggestions for figure plotting in Python. After using this tool, the authors reviewed and edited the content as needed and take full responsibility for the content of the published article.

## Author Contributions

FDB, ME conceived and designed the study. ME, ASh, AK, ASk performed the experiments. ME, ASh, FDB analyzed the data. ME, ASh, FDB wrote the manuscript. All authors discussed the results and contributed to the final manuscript. FDB: Conceptualization, Project administration, Supervision. ME, ASh, AK: Methodology, Investigation. ME, ASh: Formal analysis, Data curation. ME, FDB: Writing – original draft. All authors have read and agreed to the published version of the manuscript.

## Acknowledgments

This manuscript was developed under the project “Yeast-based biosensors for the specific and accessible detection of pathogens and antimicrobial resistance” (Acronym: AntiRYB, Reference Number: JPIAMR_2019_P08) from the Antimicrobial Resistance Joint Programming Initiative (JPI AMR) and the Research Council of Norway (RCN project number 311248). The work performed at the National Medicines Institute was funded by the National Science Centre, Poland under JPI-EC-AMR Call 2019 (NCN project number 2019/01/Y/NZ6/00006). MS-data is handled under Notur/NorStore Project NN9036K/NS9036K by PROMEC, a member of the National Network of Advanced Proteomics Infrastructure (NAPI), which is funded by the RCN INFRASTRUKTUR-program (295910).

## Data Availability Statement

The datasets generated for this study can be found via ProteomeXchange with identifier PXD055744 (Username: Password: XNtCwVbI4AIs).

## Supplementary Material

The supplementary material includes the following tables, which are provided as XLSX files:

Supplementary Table S1: Dose-correlation statistics for control strain.

Supplementary Table S2: Dose-correlation statistics for clinical strain.

Supplementary Table S3: Tabular GO-term enrichment results corresponding to **Figure 5b**.

Supplementary Table S4: Tabular GO-term enrichment results corresponding to **Figure 5c**.

Supplementary Table S5: Tabular GO-term enrichment results corresponding to **Figure 6b**.

Supplementary Table S6: List of protein IDs associated with GO terms in **Table 5**.

